# Network reaction norms: taking into account network position and network plasticity in response to environmental change

**DOI:** 10.1101/705392

**Authors:** Tyler R. Bonnell, Chloé Vilette, S. Peter Henzi, Louise Barrett

## Abstract

Recent studies have highlighted the link between consistent inter-individual differences in behaviour and consistency in social network position. There is also evidence that network structures can show temporal dynamics, suggesting that consistency in social network position across time does not preclude some form of plasticity in response to environmental variation. To better consider variation in network position and plasticity simultaneously we introduce the network reaction norm (NRN) approach. As an illustrative example, we used behavioural data on chacma baboons, collected over a period of seven years, to construct a time series of networks, using a moving window. Applying an NRN approach with these data, we found that most of the variation in network centrality could be explained by inter-individual differences in mean centrality. There was also evidence, however, for individual differences in network plasticity. These differences suggest that environmental conditions may influence which individuals are most central, i.e., they lead to an individual x environment interaction. We suggest that expanding from measures of repeatability in social networks to network reaction norms can provide a more temporally nuanced way to investigate social phenotypes within groups, and lead to a better understanding of the development and maintenance of individual variation in social behaviour.

## Introduction

Consistent inter-individual differences in behaviour are common in animal groups [1, 2]. How such behavioural differences develop, are maintained, and selected form the focus of current research efforts, as do questions relating to the ecological and evolutionary consequences of such inter-individual differences [3, 4]. There is also great interest in the extent to which animals are able to vary their behaviour in response to environmental changes, and whether and how this behavioural plasticity co-varies with inter-individual differences in mean behavioural rates. This is captured by the notion of a “behavioural reaction norm” (BRN)—the set of behavioural phenotypes an individual produces in a given set of environments—a concept drawn from life history theory [5, 6]. Analytically, a BRN can be identified using multi-level regression models, where the fitting of random intercepts captures inter-individual differences in the expression of a given behaviour, and random slopes capture individual differences in plasticity across an environmental gradient. Thus, BRNs provide information on how animals differ both in their mean level of a given behavioural trait, and in how strongly they respond to environmental variation.

Within social groups, patterns of inter-individual consistency have also been observed in social network position (e.g., central individuals remain central across time) [7-11]. Again, it is an open question as to how these differences arise and are maintained [12]. There is also evidence to suggest that consistent inter-individual differences in behaviour and social network position interact [13, 14], which implies either that, for social animals, network position reflects certain behavioural predispositions, and/or that certain behavioural predispositions may arise in response to occupying a particular network position.

At the same time, there are data showing that network structure itself exhibits strong temporal dynamics. This suggests that temporal consistency in social network position does not preclude some form of plasticity in response to environmental variation, and that network position and plasticity may co-vary in interesting ways [15]. Such co-variation could provide insight into the possible constraints or opportunities faced by individuals in the group. For example, a constraint might occur if individuals occupying more central positions in the network were in some way limited in how they could respond to environmental change. Conversely, differential opportunities might occur if certain network positions afford beneficial options with respect to coping with environmental changes in ways that were not available to others. Examining the covariation between network position and plasticity may act as a guide to discovery of the mechanisms that produce, and functional consequences of, social network structures.

Here, we use a behavioural reaction norm approach [5] to model inter- and within-individual changes in social network position—a “network reaction norm” (NRN)—and use this to quantify individual variation in social phenotypes within a troop of wild baboons as a proof-of-concept. We use a social network constructed from observed proximity data to investigate (i) whether some animals are more consistently central – measured as eigenvector centrality – over time (i.e., inter-individual differences in intercepts); (ii) the extent to which individuals on average respond to social and environmental changes (i.e., mean network plasticity in the population/group); (iii) the extent to which individuals differ from this mean response (i.e., inter-individual differences in network plasticity or variation in slope); and finally, (iv) to quantify the relationship between inter-individual differences in mean network position and network plasticity (i.e., covariation between intercepts and slopes), e.g., do more (or less) central individuals display a stronger (or weaker) response to environmental changes?

## Methods

### Networks through time

Social networks are often built on data aggregated over a specific time period. To gain an understanding of how social network structures vary through time, the most common approach has been to generate a sequence of individual time-aggregated networks and analyse the differences between them [e.g., 16]. That is, one generates a series of static “snapshots” of the social network. By necessity, this approach cannot assess any changes in the relationships that occur within the period used to construct the network, and information is therefore lost on the dynamics of social behaviour that may prove relevant and important. To address this, we make use of a moving window design, where a window of a fixed duration (anything from hours to months, depending on the question of interest—see below) shifts by a set amount of time that is smaller than the size of the window, with networks constructed at each shift [17]. This generates a time-series of networks, where the extent of the shift determines how much data overlap there is between networks. By varying the amount of overlap and the size of the window, it is possible to control the resolution at which changes in the social network are measured. This makes it possible to examine how social networks change at various levels, e.g., at the dyadic, ego-centric, or whole (sociocentric) network level.

### Individual network centrality

To investigate social network structure through time, we used proximity data collected between June 1997 to October 2006 from all adult females of a troop of baboons in the De Hoop Nature Reserve, South Africa [15, 18]. Data were collected by scan sampling every 30 min, during which the identities of all animals within 10m of the target individual were recorded (hereafter nearest neighbours) for a total of 13,504 scans across the study period. Three animals with fewer than 500 observed nearest neighbour events were removed from the sample. Given the goal of assessing individual differences in response to environmental change, this removal highlights our focus on within-individual observations [19]. The final dataset consisted of 28 females, where the median number of females at any given time was 12 (min = 9, max = 16) over the entire study period.

In selecting the temporal scale of the study, we used a window size of four months, following Henzi et al. [15] for the same population and used the netTS package in R [17]. To ensure the networks were comparable through time, we divided the observed number of nearest neighbour events by the total sampling time in the field within each window. This produced edges in the network that correspond to rates of observed proximity behaviour between any two individuals. Similarly, to ensure that the network measures were robust, we excluded networks constructed using fewer than 160h of data collection and where the correlation between the observed network measures and network measures generated from bootstrapped samples was below 0.99, which resulted in three gaps in our time series. We also removed individual measures that extrapolated beyond the range where individuals were observed (if individual A was first observed on day 100, and died on day 200, network measures for this individual were removed when the individual was only partially present, e.g., network measures for this individual generated by a window between day 80 and 120 would be removed). Using this time series of networks we extracted individual measures of eigenvector centrality (EC) [20], producing a time series of centrality measures for each individual. EC assesses the centrality of each node based on their direct and indirect connections to others. So, for example, in a fully connected network all nodes receive equal centrality scores, whereas at the other extreme, a network where all dyadic ties involve one particular individual (e.g., a star network) will result in this individual receiving the highest centrality score. More generally, the use of centrality measures that take into account indirect connections has proven useful in better understanding the fitness and heritable nature of network positions within social groups [21].

### Quantifying the changing social and physical environment

To estimate how individual social behaviour within the troop responded to environmental changes, we used two ecological variables and two social variables. We used solar radiation, measured as Watts/m^2^, at the leading edge of the moving window, as an environmental effect with the potential for immediate impact on thermoregulatory behaviour [22-24]. Estimates of surface solar radiation for our study site and study period were acquired from the Heliosat dataset, which uses the Heliosat algorithm and geostationary satellites to estimate solar surface irradiance at a spatial resolution of 0.05° × 0.05° [25]. We used cumulative rainfall within each window as an index of the amount of available food at the site [26, 27]. As such, cumulative rainfall represents an environmental variable on a coarser time scale compared to the immediate changes in solar radiation. These two environmental measures showed a low correlation (r = - 0.20; 95% credible interval (CI): −0.23, −0.17).

To capture social changes in the group across our time-windows, we used a measure of inter-individual encounter rate, and the number of males in the group. Encounter rate was constructed by using the total number of encounters observed within a time window divided by the time spent observing, e.g., if, during 100 hours of observation, there were 50 instances recorded of individuals being within 10m of each other, this would correspond to an encounter rate of 0.5 encounters/hour. This measure thus provides a group-level estimate of the rate of spatial interactions between individuals, but does not contain information on how these encounters structure at the network level. Encounter rate was moderately correlated with rainfall (r=0.44; 95%CI 0.43, 0.45), and minimally with solar radiation (r=−0.15; 95%CI: −0.16,-0.14). The number of males was a count of adult males present in the group during any given time window of observation. Female social relationships, in this case measured through proximity, are thought to be influenced by the number of males in the group through increased/decreased female competition for infant protection from infanticide by associating with available males [28]. Male number was obtained from demographic records maintained on the group across the entire study period.

### Network reaction norm

To model individual eigenvector centrality, we used both random intercepts and slopes in a multilevel model, and interpret our model within a behavioural reaction norm framework [5, 6] (Fig. 1). In this framework, random intercepts indicate inter-individual differences in mean network position, while random slopes indicate individual differences in network plasticity (Fig. 1).

**Figure 1:**
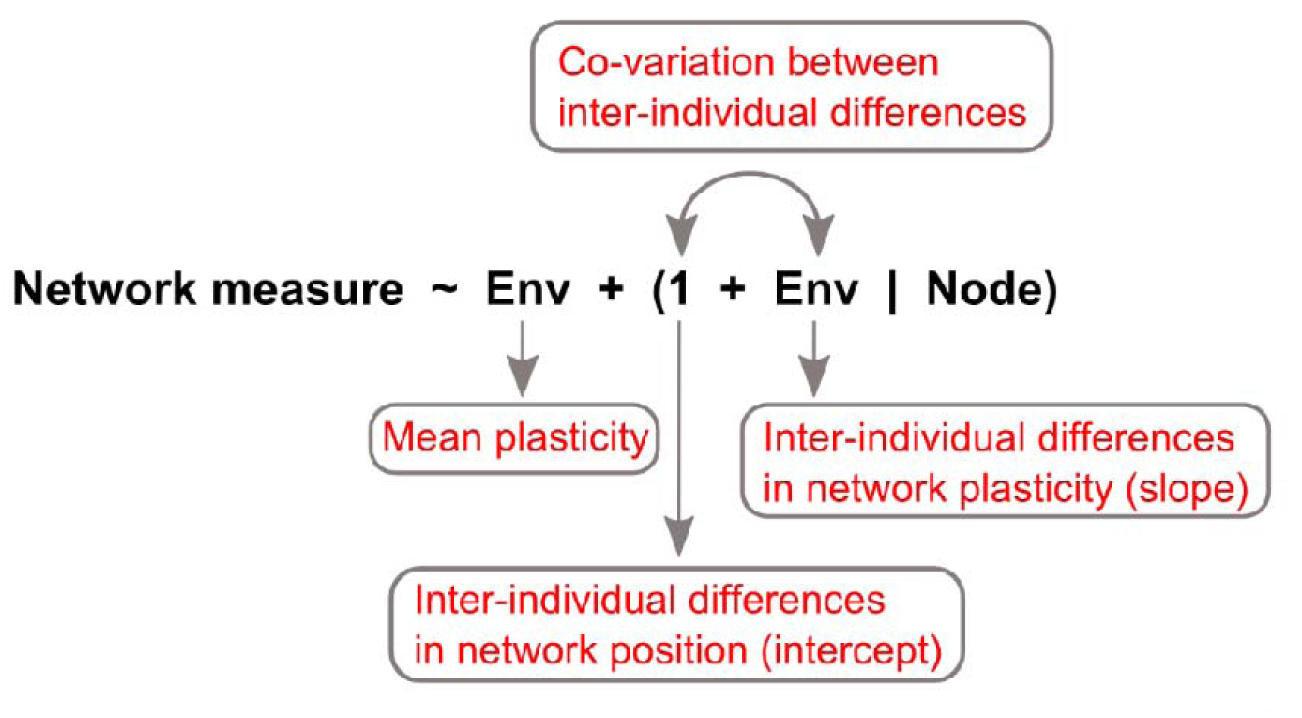
Pseudocode for the interpretation of a multilevel model using the network reaction norm (NRN) framework. The format of the pseudocode follows that of multilevel model specifications in the R programing language. See Table S1 for further details.

Given the time series nature of the data, we used generalized additive mixed models (GAMM) to allow for non-linear relationships, and use penalized regression splines to avoid overfitting [29]. To account for potential sampling bias, e.g., the possibility that some individuals were sampled only in high rainfall conditions, we included the mean environmental conditions experienced by each individual. This allowed us to partition within- and between-individual effects of the environment [30]. We allowed the individual effects of environmental conditions to be non-linear by including a smooth term, using thin plate regression splines, and allowed all individuals to vary in their response to environmental changes by including random slope terms for each environmental variable.

We also included the possibility of a non-linear seasonal trend in centrality values by using day of year and a cyclic cubic regression spline to account for the circular nature of this variable. Rather than stripping autocorrelation from the data we accounted for it in the residuals using an ARMA(1,1) process [31]. We used the R package “brms” [32] to fit the model in a Bayesian framework. We scaled and centred our independent variables, and used weakly-informative priors centered on zero, i.e., Normal(0,1). This reflects our prior assumption of no effect, and reduces problems of potential collinearity between moderately correlated independent variables [33]. Model diagnostics suggest MCMC convergence, with all R□<1.05. All analyses were run in the R programming environment 3.5.2 [34].

### Predicting network positions in changing environments

While one use of behavioural reaction norms is to quantify variability in network position and plasticity within a population, it can also be used to predict how network positions will change as environments change. To highlight this, we withheld approximately one year of data (i.e., the final period of data collection between 2005-08-15 to 2006-10-15), and used this to assess the ability of the fitted model to make accurate predictions about network positions on novel data. This also helps assess whether the model is overfitting, and allows us to measure the importance of individual-level effects, i.e., does individual identity help to predict changes in network position outside of the data used to fit the model? More importantly, by testing the model on a new dataset, we can determine how well the observed effects of social and physical environmental changes generalize to new conditions. To do this, we first estimated the amount of variation (R^2^) in individual eigenvector centrality that our model could explain in the withheld dataset, using only those individuals found in both datasets. We then took advantage of two time periods in the withheld dataset where solar radiation varied from high to low (317 to 103 W/m^2^). Making predictions about changes in individual network position during these two time periods allowed us to test whether the predicted changes in centrality match those actually observed.

## Results

### Network reaction norms

In the case of behavioural reaction norms, the parameters of interest are the inter-individual differences (i.e., random effects). We therefore proceed by presenting the inter-individual differences in mean network position, then present the results of mean network plasticity and inter-individual differences in this plasticity, followed by the covariation between inter-individual differences in network position and plasticity (i.e., the correlations between random intercepts and slopes). Finally, we present estimated reaction norms for each individual across our environmental gradients to assess the stability in the rank order with respect to occupation of central positions in the group.

The fitted NRN suggests there are large inter-individual differences in network position, identifying mean centrality as the parameter with the largest inter-individual differences (sd(μ) = 1.16, Table 1, Fig. 2). The “sd” here refers to the standard deviation of the individual differences, a hyper-parameter estimated in the model that determines the magnitude by which individuals differ in their estimates. The NRN also suggests that individuals exhibited network plasticity, i.e., that changes in network position were related to changes in environmental conditions.

**Table 1:**
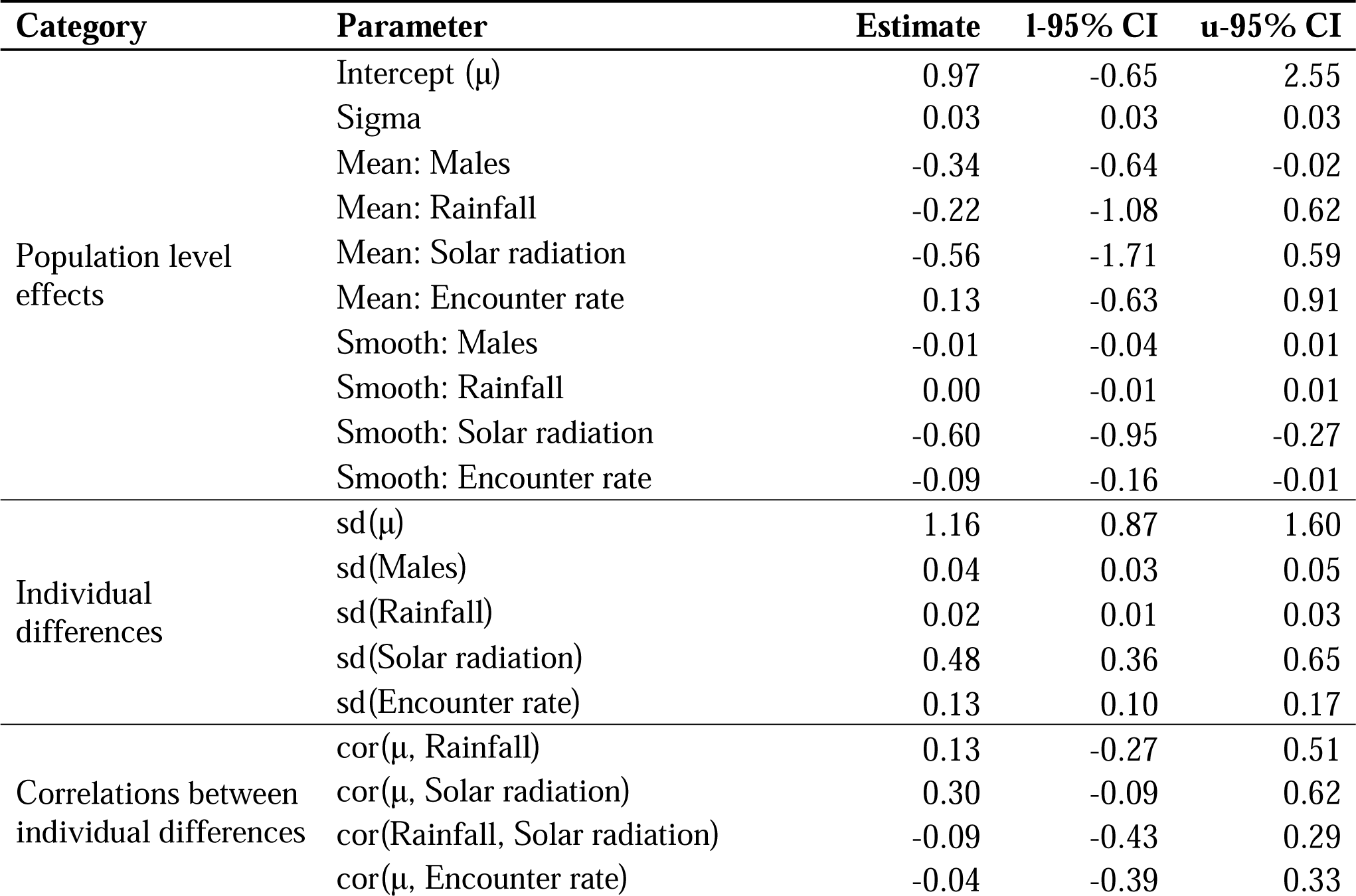
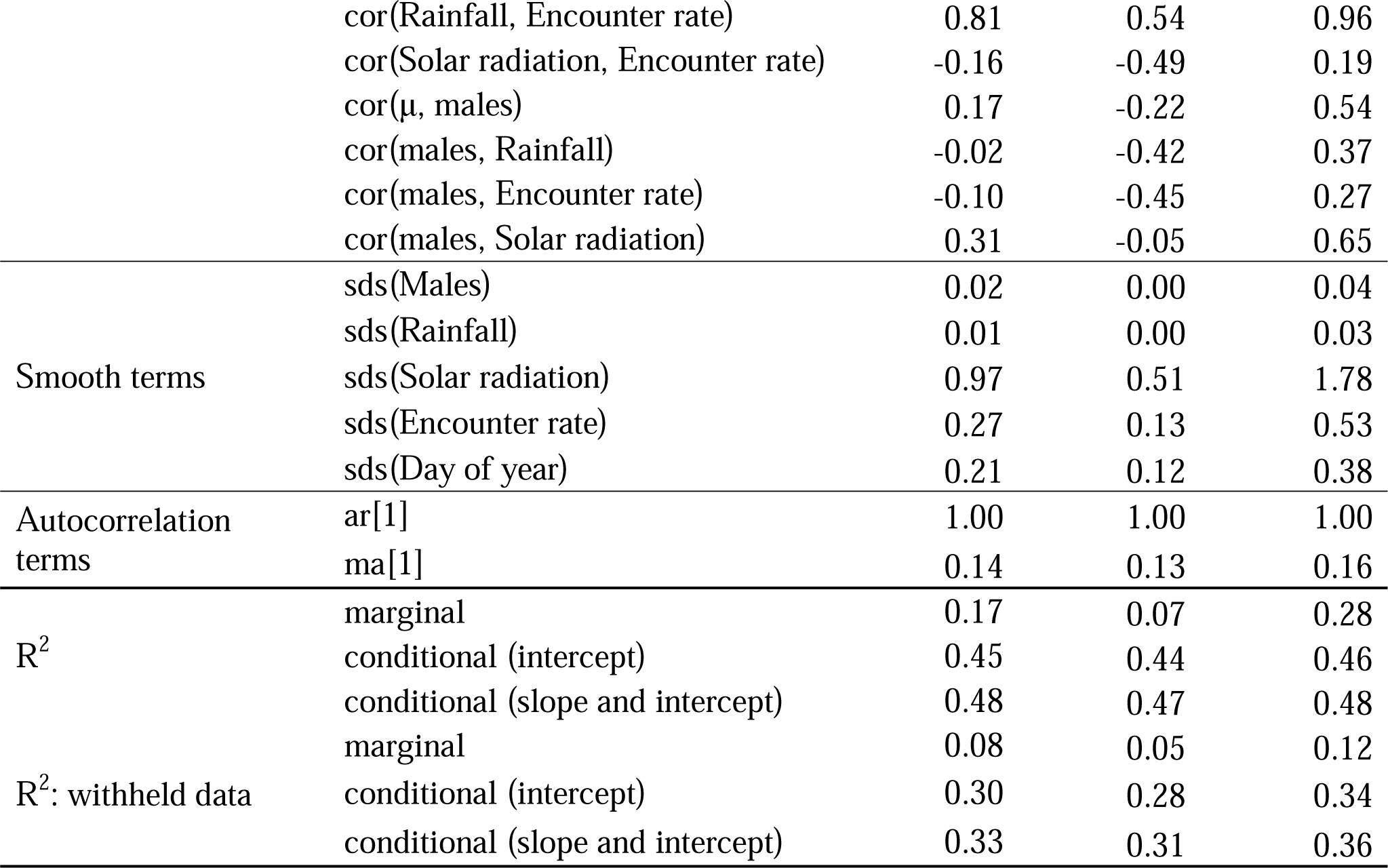
Parameter estimates (±95% CI) for the NRN. We present the parameter estimates of the GAMM partitioned into 1) the population level effects containing the mean responses across the population, 2) the magnitude of individual differences in responses (random effects), 3) the estimated correlation between individual differences, and 4) the magnitude of the complexity of the smooths, controlled by a smoothness term (sds) where larger values indicates a more complex fit. Given the use of regression splines to estimate the relationships between independent and depended variables, the table is accompanied by plots (Figs 3,4), as the interpretation of these relationships are not possible from the table alone. The autocorrelation terms ar[1] and ma[1] are respectively the estimated magnitude of the autoregressive process of order 1, and the moving average of order 1. Finally, both the in- and out-of-sample R^2^s are presented for both main effects (marginal) and with random effects (conditional) included.

**Figure 2:**
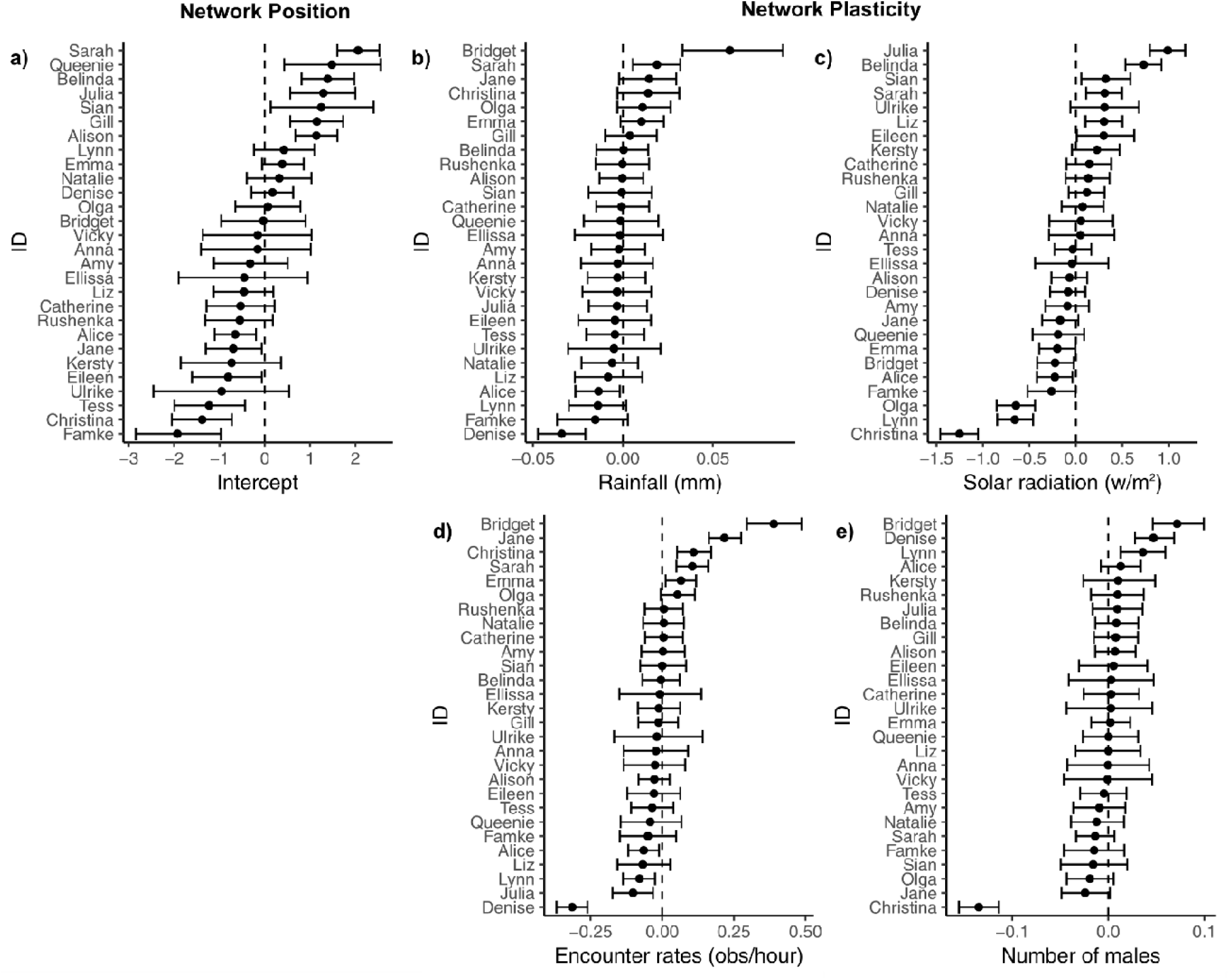
Estimated inter-individual differences in a) mean network centrality (i.e., random intercepts), b-e) response to environmental changes (i.e., random slopes). The vertical line at zero represents the mean, and points (±95% CI) indicate individual deviation from the population mean. Individual names on the y-axis where ordered highest to lowest in each plot to highlight deviations from the mean intercept (a) and deviation from the mean slope (b-e) (i.e., 0).

With respect to physical environmental changes, we found a mean response to changes in solar radiation, but none for rainfall (Table 1, Fig. 3). On average, individuals decreased in eigenvector centrality as solar radiation increased. For changes in the social environment, the population showed no mean response either to changes in encounter rate or to the number of males in the group (Table 1, Fig. 2). There was also a mean seasonal response in network positions, with individuals on average having lowest centrality values in June-July (which is the Austral winter) (Table 1, Fig. 2).

**Figure 3:**
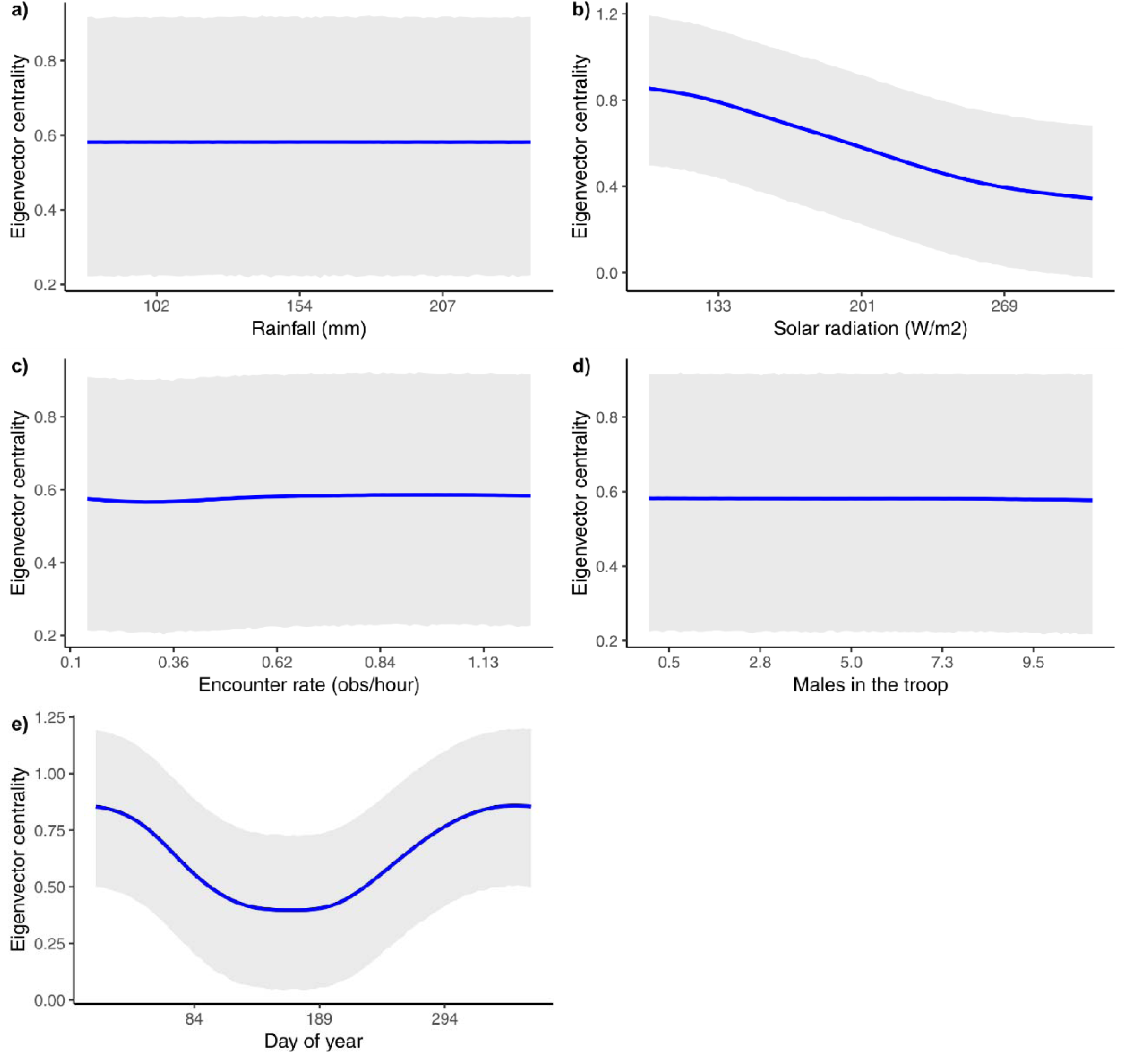
Mean response of network position, in terms of individual eigenvector centrality, to changes in physical and social environment: a) cumulative rainfall, b) solar radiation, and c) group encounter rates, d) number of males in the troop, and d) day of year. The mean estimate (blue line) and 95% credible intervals (shaded areas) are presented for each.

Individuals also differed in mean network plasticity, with the largest differences coming from responses to solar radiation (sd(Solar) = 0.48) and encounter rates (sd(EncounterRate) = 0.13), with little evidence for any inter-individual differences in response to changes in rainfall, or number of males in the troop (Table 1, Fig. 2).

We found no strong evidence for co-variation between inter-individual differences in both network positions and network plasticity. Although there was a correlation between individual difference in response to rainfall and encounter rates, the magnitude of the individual differences themselves suggest a very weak effect at best, and caution is needed in interpreting this result (Table 1, Fig. 2).

Overall, the NRN suggests a substantial amount of variability can be explained by individual differences in network position and plasticity; using R^2^ as a measure of explained variance [35], the main effects accounted for 0.17% of the variation in the data (i.e., Marginal R^2^), whereas taking into account the individually-varying intercepts and slopes explained 48% (Conditional R^2^) (Table 1).

To address the question of whether some individuals were more consistently central than others, we plotted reaction norms against changes in each environmental variable (Fig. 4). The results suggest that rankings of individual centrality values are influenced by solar radiation and, to a lesser extent, encounter rate and number of males, but not by rainfall.

**Figure 4:**
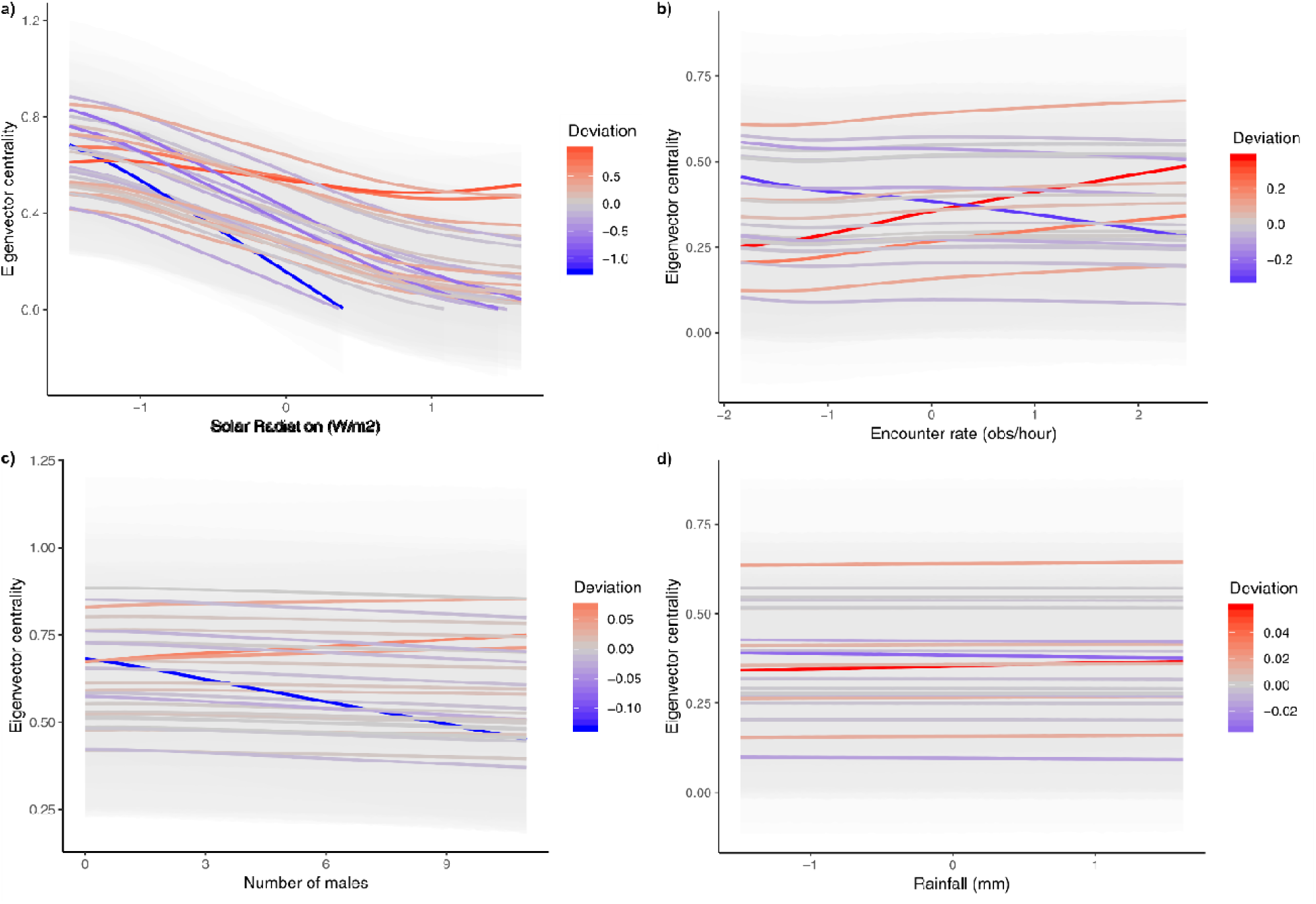
Reaction norm plots of individual centrality in response to changes in a) solar radiation, b) encounter rates observed in the group, c) number of males in the group, and d) rainfall. Blue lines identify individuals with slopes that are lower than the average for the group, and red line indicate those with slopes that are higher than the average for the group.

### Predicted network positions in the withheld dataset

When comparing how individuals changed in network positions between the two time periods in the withheld dataset (Fig. 5a dashed lines), we found that the predicted relative changes in an individual’s eigenvector centrality (i.e., whether centrality increased or decreased qualitatively) did not closely match observed changes (Fig. 5b). Similarly, when we look at individual eignvector centrality values in absolute terms (i.e., whether there was a good quantitative fit), we found that many individuals did not match observed levels (Fig. 6cd, where flat lines indicate a match between predicted and observed, and slopes indicate a mismatch), although it should also be noted that predictions were better for some individuals than others. This difference in predictability might suggest that some individuals have more predictable responses in network positions compared to others, and that this within-individual variation might usefully be explored further within the reaction norm framework [6]. Additionally, when we considered variation in eigenvector centrality at the population level, the model was able to explain 33% of the variance in the withheld dataset, suggesting that the predicted relative differences in eigenvector centrality do hold some predictive capacity (Table 1).

**Figure 5:**
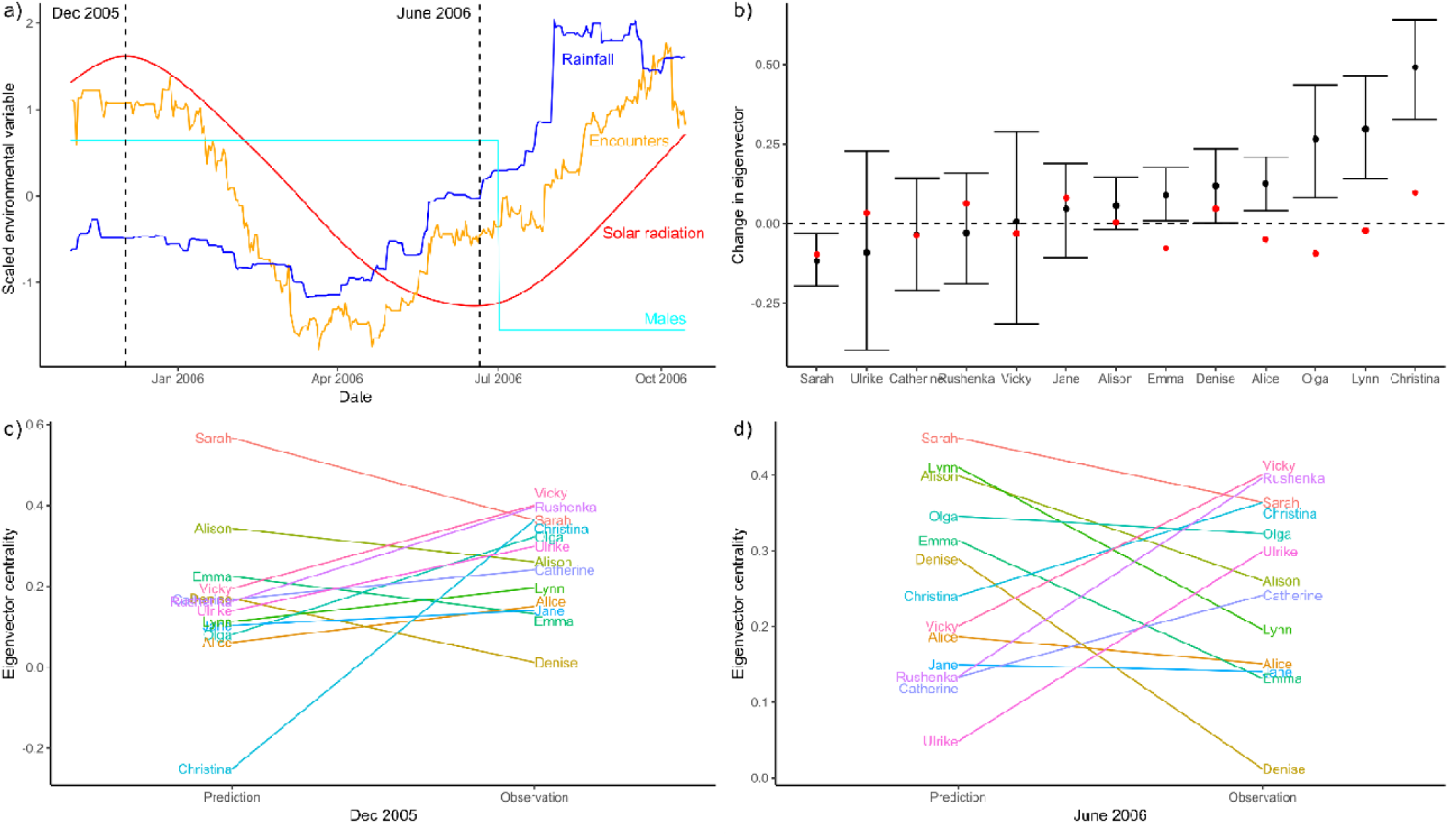
Withheld data, not used in the model fitting process, is used to test how well the model generalizes to novel environments. (a) Solar radiation, rainfall and encounter rates measured across the withheld dataset. The dotted lines indicate the two time points used to make predictions: maximum solar radiation (Dec 2005) and minimum solar radiation (June 2006). (b) Predicted changes in network position between Dec 2005 and June 2006. Black circles represent predicted centrality changes, with error bars representing 95% credible intervals. Red dots indicate observed changes in individual centrality. Predicted vs. observed eigenvector centrality values for each individual for (c) December 2005 and (d) June 2006. Lines are drawn between predicted and observed points to facilitate comparison, i.e., flat line indicates that prediction matches observation. To aid in interpretation of the changes in eigenvector centrality (y-axis), the range of eigenvector centrality values in the observed data was between 0 and 0.54.

## Discussion

Overall, the use of the network reaction norm approach suggests that variation in eigenvector centrality in our study group is driven largely by consistent inter-individual differences in mean network position, where some individuals are, on average, higher in centrality than others. Beyond inter-individual differences in mean centrality, we found evidence of a mean network plasticity to changes in only one environmental variable (solar radiation) suggesting a similar group wide response. Within-individual differences were also found in network plasticity to solar radiation, encounter rates, and number of males in the troop, suggesting that a given individual’s network positions did not respond in the same way as others to changes in their social and physical environments. These within-individual differences were, however, estimated to be much smaller than inter-individual differences in mean centrality (Table 1). A comparison of differences in network position and plasticity also showed very little co-variation, suggesting that there were no strong constraints or opportunities faced by individuals due to their network position, e.g., more central individuals did not show increased/decreased network plasticity in response to environmental change.

Given that our estimated NRN indicates that some individuals occupied network positions that were more/less responsive to changes in the environment, it raises the question of what kind of mechanisms are driving this variation: is it the case that individual differences in genetic traits and/or developmental environments are responsible for this variation in network reactivity [36]? Or is there something about network position beyond eigenvector centrality that underpins more/less reactive responses? This variation in network plasticity also suggests consequences for who is likely to be the most central animal in the group, and under what environmental conditions. For example, individual rank order in terms of who is most central in the group is predicted to differ under low, compared to high, solar radiation conditions (Fig. 4a). More generally, the presence of a network position x environment interaction carries implications for estimating the fitness-related benefits that individuals acquire from particularly advantageous positions within a social network. For instance, if there are beneficial network positions within a population [37-40], the presence of individual differences in network plasticity suggests that environmental conditions can have a direct impact on which individuals will be found in beneficial network positions and over what period of time. This suggests that when relating individual fitness outcomes to social network positions a more dynamic approach that includes changes in the environment will be necessary when the population shows network position x environment interactions.

The ability to quantify individual variation in network position and plasticity facilitates various forms of comparisons across species, or within species across populations, to provide insights into the evolution and development of social behaviour more generally. For example, if we consider comparisons within-species and across different study sites, it is possible to investigate whether individual variation in centrality is largely due to inter-individual differences in mean centrality (i.e., intercepts)—as our study suggests—or determine whether some populations also show greater individual differences in network plasticity and, if so, under what environmental conditions. When comparing across species, one could investigate whether phylogenetic relatedness and/or environmental conditions predict the magnitude of inter-individual differences in mean network position and/or inter-individual differences in network plasticity. Additionally, by combining NRNs and BRNs it should be possible to look at both adaptability in social structuring (NRN) and social behaviour (BRN), quantifying flexibility in social structure (e.g., whether and how overall network structure changes) and flexibility in social behaviour (i.e., how individuals vary their behaviour in order compensate for environmental changes, and so preserve network/social structure) [34]. This combined approach might be very useful when investigating animal responses to (rapidly) changing environments. This is generally predicted to be the case for primate social groups (e.g., vervet monkeys, *Chlorocebus spp*.), while for other species, such as the African striped mouse (*Rhabdomys pumilio*), changes in environment lead to drastic changes in social organization with little change in individual behaviour [15, 36, 41].

The use of a withheld dataset with NRNs can expand beyond the quantification of social variability in a population, to allow for tests of how network position x environment interactions generalize to future conditions. In our case the withheld dataset confirmed that the general pattern in individual variability held, with most variation in centrality (R^2^) being explained by inter-individual differences in intercepts. However, the results from the withheld dataset suggest that we have poor predictive capacity with respect to shifts in centrality in response to future environmental conditions. This suggests either that (a) changes in eigenvector centrality are, in large part, stochastic, (b) that any new individuals entering the testing dataset are highly influential, or (c) that we have not found the relevant social and environmental drivers of individual centrality. The use of a withheld dataset is therefore a particularly useful tool for identifying and testing these possibilities, and making assessments of the predictive component of NRNs.

It is also important to recognize that a population’s rate of turnover, and the strength of inter-individual differences in a group (and population), will set limits to the temporal scale for predicting social network changes using the NRN approach [42]. If inter-individual differences in network position and plasticity are found to be important then, as long as the same individuals are present and turnover in the population is low, there is likely to be high predictability across time. Conversely, when population turnover is high, predictability will drop as there will be no available estimates of inter-individual differences from past data. However, in cases where there are no strong inter-individual differences in network position and plasticity, and mean network plasticity is strong (i.e., all animals respond in the same way), then predictions are likely to be viable over longer time scales.

More generally, the ability to make predictions about social networks in changing environments can have direct implications for disease management, population level dynamics, as well as other ecological and evolutionary processes where social network structures have been shown to be an influential component [43]. Additionally, increases in geospatial infrastructure have made a growing amount of environmental data available, particularly from freely available satellite remote sensing datasets (e.g., Sentinel-2, Landsat, Modis) [44, 45], facilitating the integration of dynamic social networks with environmental measures. This suggests that a NRN approach could be a very useful approach for studying the response of social species to both landscape land use changes or climatic changes.

By shifting towards a dynamic view of social networks it is possible to better understand socio-structural changes in animal groups. We have demonstrated that a NRN framework with a dynamic social networks approach, makes it possible to quantify individual variability in both network position and network plasticity. The ability to consider individual network positions and environmental contexts simultaneously also serves to facilitate the interrogation of their interaction.

## Acknowledgements

We thank Cape Nature for providing permission for the baboon research, and a host of students and assistants who contributed to the long-term data set.

## Electronic Supplementary Material

**Table S1:** We present a multilevel model for a single network measure and a single environmental variable (Env), assuming a longitudinal (i.e., time-series) dataset. The network reaction norm interpretation is presented alongside the multilevel model formulation. We use mathematical syntax following Stan (Stan Development Team 2018).

| Equation/Parameters | Multilevel model | NRN interpretation |
| --- | --- | --- |
| $Network\_measure_{i,t} \sim Normal(mu_{i,t}, \sigma_{\epsilon})$ | Likelihood | - |
| $mu_{i,t} = \alpha + \alpha_i + (\beta + \beta_i) * Env_t$ | Linear model | Mean network measure of node $i$ at time $t$ |
| $\alpha$ | Mean intercept | Mean network position |
| $\beta$ | Mean slope | Mean network plasticity |
| $\alpha_i \sim Normal(0, \sigma_{\alpha})$ | Random intercepts | Individual differences in network position |
| $\beta_i \sim Normal(0, \sigma_{\beta})$ | Random slopes | Individual differences in network plasticity |
| $\sigma_{\alpha}$ | Standard deviation of the differences in intercepts | Magnitude of differences in network position |
| $\sigma_{\beta}$ | Standard deviation of the differences in slopes | Magnitude of differences in network position |
| $\sigma_{\epsilon}$ | Model standard deviation | Uncertainty in predicted network position |
| $corr(\alpha_i, \beta_i)$ | Dependence between random intercepts and slopes | Dependence between network position and plasticity |

